# Aminoglycosides induce a bacterial senescent state that increases antibiotic tolerance in treatment-naïve cells

**DOI:** 10.1101/2021.10.04.463054

**Authors:** Christian T. Meyer, Giancarlo N. Bruni, Ben Dodd, Joel M. Kralj

**Author notes:** UCLA, Department of Genetics. University of Michigan, Department of Human Genetics (Ben Dodd). These authors contributed equally (Christian T. Meyer, Giancarlo N. Bruni, Ben Dodd).

## Abstract

Bacterial evolution of antibiotic resistance is facilitated by non-genetic resistance that increases drug tolerance, buying time for evolutionary innovation. *Escherichia coli* treated with aminoglycosides permanently lose the ability to divide within four hours, yet we discovered a majority of cells maintain membrane integrity and metabolic activity greater than two days post treatment - a bacterial senescent-like state. These cells, which we term zombies, exhibit dynamic gene expression and metabolomic profiles, even after irreversible exit from the cell cycle. Our data reveal zombies upregulate the phage shock protein pathway to maintain membrane integrity. Remarkably, though unable to form new colonies, zombies increase the antibiotic tolerance of treatment-naïve cells, implying chemical communication. Chemical supplementation and genetic knockouts show that zombies communicate with treatment-naïve cells by secreting indole. In summary, our study revealed a bacterial senescent-like state, induced by aminoglycosides, that decreases the antibiotic susceptibility of multiple bacterial species. Thus, *E. coli* zombies utilize paracrine signaling to promote non-genetic antibiotic tolerance.

## 2. Introduction

Effective antibiotics are a pillar of modern medicine, albeit one that is rapidly losing utility. Bacteria continually develop new methods to survive lethal antibiotic treatments, commonly via genetic resistance. However, it has been shown that antibiotic tolerance, the ability of cells to survive a lethal treatment in the absence of a heritable resistance, precedes the formation of genetic resistance^1^. Maintaining the current era of antibiotic supremacy will require a deeper understanding of the cellular and population response to current drugs^2,3^. Better understanding will enable new compounds and therapeutic strategies to target bacteria exhibiting non-genetic resistance^4–7^.

Non-genetic resistance is commonly considered a property of individual cells. However, bacteria most often live within mixed communities, constantly adapting to each other and environmental stimuli via paracrine signaling. Communities of cells often communicate to coordinate and diversify the population level response to changes in environment^8–10^. Intra- and inter-species communication is the basis for both competition and cooperation, and a sophisticated chemical lexicon has evolved within bacterial communities^11–13^. Therefore, paracrine signaling represents a potential source of non-genetic resistance arising only in the context of bacterial communities. It is necessary to develop a population-level perspective that integrates outcomes at the scale of individual cells, and how those outcomes propagate to the emergent behavior of the bacterial community. Such a perspective will allow researchers to take a more nuanced approach to evaluate and understand our current therapies and how bacterial communities evade them.

For over 100 years, colony forming units (CFUs) have been the gold standard to measure life and death of bacteria, relying on the growth of a bacterium to form a mass large enough to be visually detected. Technological advances such as plate readers can query thousands of conditions but still rely on bulk population measurements. Despite the power, utility, and success of these assays, they are necessarily blind to the physiological changes arising in individual cells and the heterogeneity within a population. Genetically encoded fluorescent sensors have helped to shed light into this blind spot by enabling measurements with high temporal resolution at the single cell level^14–18^.

In this paper, we identified a metabolically active, non-dividing cellular state by monitoring single cell physiology upon treatment with aminoglycosides. We found that even though *E. coli* rapidly lose the ability to divide and form colonies, they continue to metabolize sugars for up to 4 days. During the time between the cessation of cell division and the cessation of metabolism, *E. coli* secrete indole and increase the antibiotic tolerance of nearby, treatment-naïve cells. In addition to promoting non-genetic resistance in neighboring *E. coli*, the secreted indole can also protect other, non-related species of bacteria. Thus, when considering the response of a bacterial community to antibiotic treatment, it is essential to consider these non-dividing cells which communicate via paracrine signaling and are invisible to traditional CFU assays.

## 3. Results

### Aminoglycoside treated cells enter a senescent-like state

Using GCaMP6f as a measure of free cytoplasmic calcium in *E. coli*, we previously found aminoglycoside treatment induced a dysregulated ionic state^19–21^. The dysregulated calcium dynamics were persistent for at least one day after exposure, despite the onset bactericidal activity of the drug occurring within 1 hour^19^. To quantify the duration of persistent, aminoglycoside-induced, calcium fluctuations, *E. coli* expressing a GCaMP6f-mScarlet fusion were imaged over 96 hours after addition of kanamycin (Fig 1A,B, Movie S1). We observed cells maintained membrane integrity as well as GCaMP6f and mScarlet fluorescence for up to 4 days. Calcium traces from individual cells showed heterogeneity in the timing of activity cessation (Fig 1B). The timing of activity cessation depended on the concentration of kanamycin (Fig 1C). At all kanamycin concentrations, the fraction of cells exhibiting calcium activity decreased over time. At 6x the minimum inhibitory concentration (MIC) of kanamycin, over 1/3 of cells maintained calcium dynamics after 72 hours of continuous treatment. Untreated cells did not show these transients and underwent normal cell division under identical imaging conditions (Fig S1). Heat killed cells, either untreated or treated with kanamycin, did not grow or divide, did not show any GCaMP6f transients, and resulted in no CFUs when plated (Fig S2A-D). In kanamycin treated cells, membrane integrity was maintained relative to heat killed cells as the magnitude of propidium iodide (PI) fluorescence influx was higher in heat killed cells (Fig S2E,F). Gentamycin, tobramycin, and streptomycin also previously showed calcium dynamics for 24 hours after CFU formation ceased^19^, which suggested this phenotype is common to all aminoglycosides. These long-term aminoglycoside-induced calcium dynamics are likely driven by non-equilibrium cellular processes, as outlined in previous work^19^. These data revealed that cells could maintain plasma membrane integrity and dynamic calcium transients for up to 4 days after bactericidal treatment.

**Figure 1:**
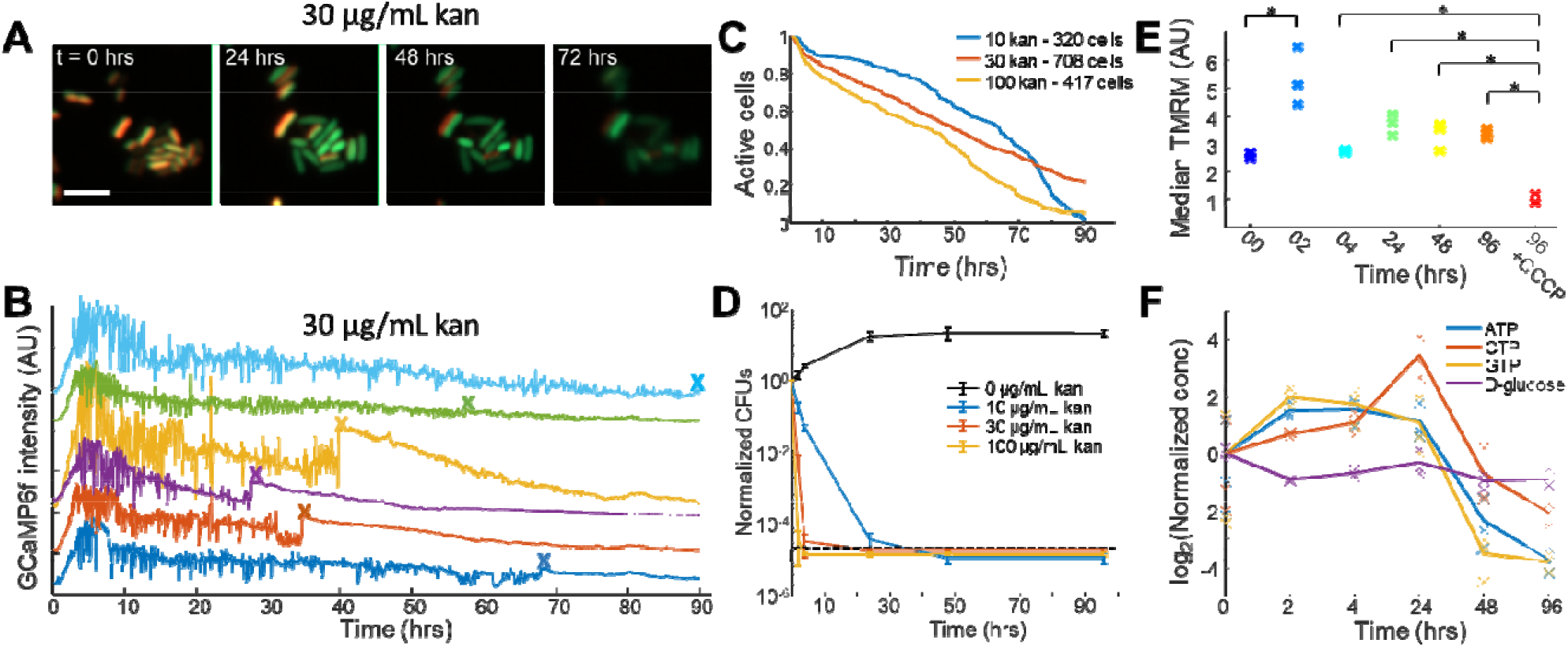
Aminoglycosides induce a senescent-like (“zombie”) state in *E. coli*. (A) Images of *E. coli* expressing GCaMP6f (green) and mScarlet (red) treated with 6x MIC (30 μg/mL) of kanamycin over 72 hours. The scale bar is 5 μm. (B) Single cell fluorescent traces from *E. coli* expressing GCaMP6f under continuous treatment of 30 μg/mL kanamycin added at t = 0 hrs. Each color is a different cell from the same field of view. The x indicates the manually defined time point of activity cessation. (C) Kaplan-Meyer curve based on the cessation of calcium dynamics. Cells were treated with 10, 30, or 100 μg/mL kanamycin at time t = 0. (D) CFU measurements upon treatment with kanamycin. Each data point is the average of 3 biological replicates, and the error bars represent the standard error. The black dashed line is the limit of assay detection. (E) Median TMRM fluorescence (proportional to membrane voltage) of 1e5 kanamycin-treated (30 μg/mL) cells measured by flow cytometry. At 96 hours, cells were also treated with 50 μM CCCP. ^*^ denotes p < 0.05 (two-sided t-test). (F) Cellular concentration of NTPs and glucose after treatment with 30 μg/mL kanamycin. Each time point had 3 biological replicates, and the line denotes the mean value of the 3 replicates.

To confirm that *E. coli* in these conditions were unable to divide, CFUs were measured across a titration of kanamycin concentrations over 96 hours (Fig 1D). As expected, kanamycin exhibited rapid bactericidal activity as a function of concentration. At 30 μg/mL kanamycin (∼6x MIC) and higher, the CFU count was below our detection limit (500 cells/mL, black line Fig 1D) within 4 hours of treatment. Antibiotic-free agar plates of kanamycin treated cells left at 30 □ for 10 days did not regrow colonies and showed the drug was indeed bactericidal, not merely bacteriostatic (Fig S3). Our CFU data are consistent with decades of previous research on the lethal outcomes of aminoglycosides, and that treated *E. coli* cannot divide and form new colonies^22–24^. In contrast, the continued calcium dynamics suggested that these cells maintain active metabolism and comprise a large fraction of the population. These data indicated that this state was distinct from persisters^25,26^ or viable but non-culturable^27^ cells that regrow after dormancy and occur in very few cells. However, the dichotomy between calcium imaging, which suggested active metabolism, and CFU data, which indicated death, encouraged us to ask if bacteria were indeed still alive after the permanently ceasing cell division.

Metabolic activity was measured across a 96-hour treatment. *E. coli* in an aerobic environment maintain a proton motive force (PMF) across the plasma membrane to power the F1Fo-ATPase^28^. We measured voltage, a component of the PMF, using the membrane-permeable dye, tetramethylrhodamine methylester (TMRM) which accumulates in cells with negative potential^19,29–31^. Upon treatment with kanamycin, *E. coli* showed an increase in TMRM uptake at 2 hours, consistent with previous observations^19,32^ (Fig 1E). With more time, the membrane potential decreased compared to the 2-hour time point, but even at 96 hours, the TMRM fluorescence was higher (more polarized) compared to untreated conditions. Treating cells after 96-hours of kanamycin exposure with carbonyl cyanide m-chlorophenyl hydrazine (CCCP), a proton uncoupler that sets the PMF to zero, decreased TMRM fluorescence (Fig 1E). Consistent with our TMRM data, the Nucleotide Tri-Phosphate (NTP) pool showed a rapid increase after treatment and was not reduced until 48 hours post treatment while internal glucose was maintained throughout (Fig 1F). Untargeted cellular metabolomics over 96 hours showed an evolving metabolic profile associated with increased amino acid uptake and fatty acid synthesis (Fig S4A-C, File S1). In order to power the electron transport chain and generate a voltage, cells must metabolize a carbon source. To check that non-dividing but polarized *E. coli* were still capable of consuming glucose, cells were treated with kanamycin for 24 or 48 hours and placed in fresh medium with kanamycin. Extracellular glucose was measured in the medium as a function of time, and we observed continued glucose depletion which demonstrated cells continued to metabolize a carbon source (Fig S4D). Thus, we concluded up to 4 days after a continuous lethal treatment of aminoglycosides, *E. coli* consumed glucose, generated ATP, established ionic gradients, and maintained membrane integrity. These cells were similar to minicells^33^ in that they maintain metabolic activity without cell division, yet they also maintain a chromosome, distinguishing them from minicells. We termed these cells ‘zombies’ as they occupied a space between the classical microbiological definitions of life and death.

### Aminoglycoside-induced zombies are maintained by the phage shock protein response

We next sought to discover mechanisms that enable continued metabolic activity in the face of bactericidal doses of aminoglycosides. To identify potential genetic pathways involved in the maintenance of the zombie state, RNA expression was measured at 6 time points up to 24 hours after aminoglycoside treatment (Fig 2A). We used RNAseq to measure gene expression changes over time after aminoglycoside treatment as it is upstream from ribosomal inhibition (File S2). A total of 1746 differentially expressed genes (DEGs) were identified at one or more time points post-treatment (File S3). Principal component analysis (PCA) showed biological replicates were well clustered, and that time points were separated along the first 2 components (Fig S5A). Early gene expression data (t = 0, 30, 90 min) matched well with previously published data sets of kanamycin treated *E. coli* (File S2)^32^. The RNAseq profile continued to evolve after 4 hours when the CFUs went to zero (Figs 2A,S5). Our data shows gene expression varied across multiple time scales with characteristic transcriptomic profiles for early and late time points, including after CFU based assays report zero survivors.

**Figure 2:**
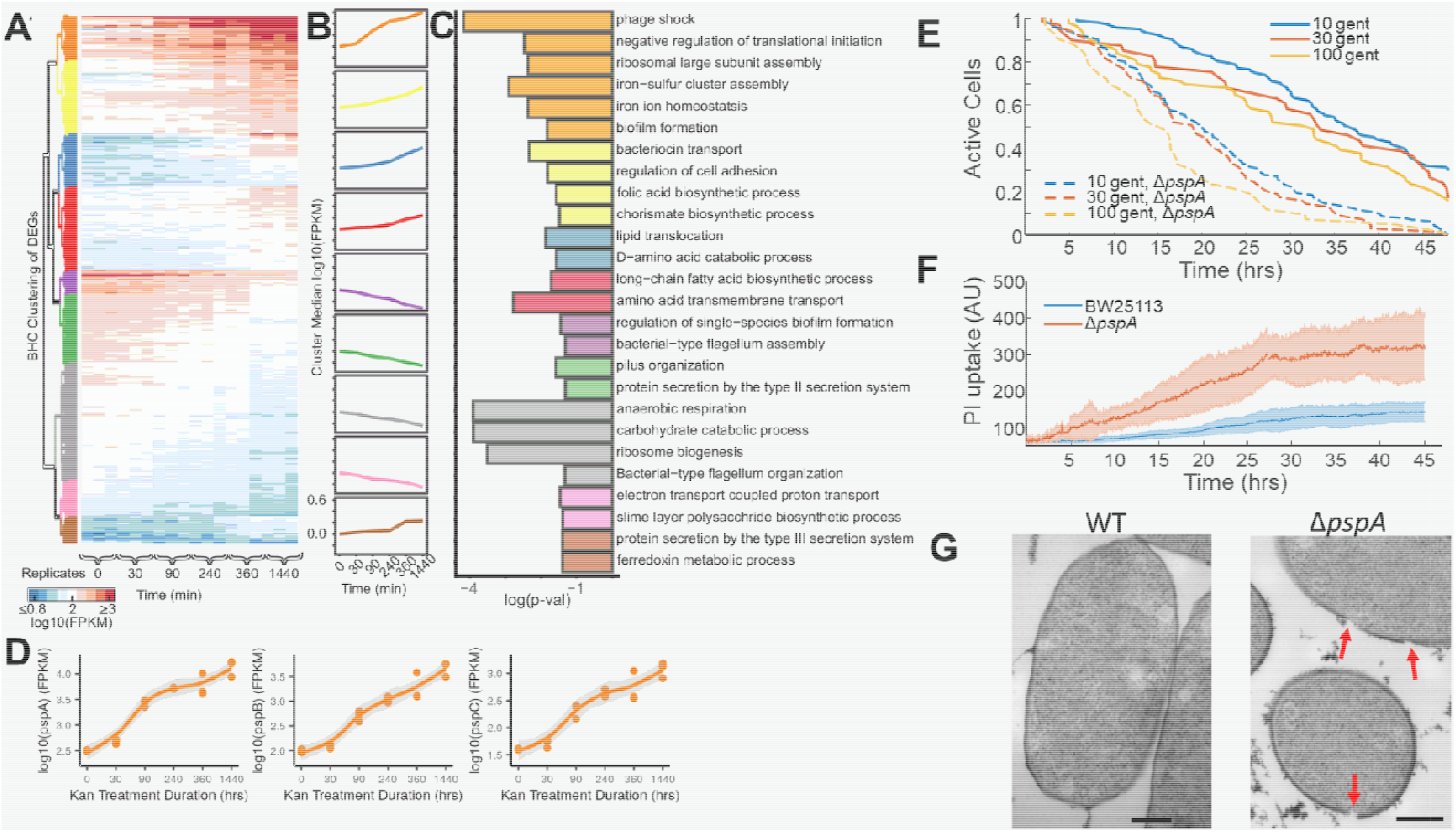
Zombie cells are maintained via phage shock protein membrane repair. (A) Heatmap of DEGs upon treatment with 30 μg/mL kanamycin. Each time point was measured in biological triplicate. Cluster (color bars left) determine using Bayesian Hierarchal Clustering (BHC) (See Methods) (B) Median expression (FPKM) of all genes in each cluster over time. (C) Enriched GO terms associated with each cluster along with the p-value (Fisher’s exact test) of over representation (x-axis). The color on the bar chart corresponds to the cluster in panel B. (D) Expression of phage-shock genes *pspA, pspB*, and *pspC* upon treatment with kanamycin. Line represents mean and shaded error bar is the standard deviation between biological replicates (points). (E) Kaplan-Meyer survival curve based on the cessation of calcium dynamics for WT cells (solid lines) and Δ*pspA* cells *(dotted lines)* upon treatment with 10, 30, or 100 μg/mL gentamicin. (F) Propidium iodide (PI) uptake for either WT (blue) of Δ*pspA* (red) upon treatment with 10 μg/mL gentamicin. The shading indicates the standard deviation across all measured cells (WT-n = 428, Δ*pspA*-n = 229). (G) TEM micrographs of WT (left) and Δ*pspA* (right) upon 4-hours treatment with 30 μg/mL gentamicin. The red arrows indicate kanamycin-induced membrane damage. The scale bars are 200 nm.

To leverage the dynamic nature of the gene expression data, DEGs were grouped using Bayesian Hierarchical Clustering^34,35^ (BHC, Fig 2A,B). Gene Ontologies (GO) enriched in a BHC cluster of genes that increased expression over time with aminoglycoside treatment were similar to previous reports, with strong enrichment in ribosome assembly, phage shock, and iron-sulfur cluster assembly (Fig 2B,C, orange cluster; File S4)^32^. Conversely, GO terms associated with decreasing expression included anaerobic respiration and electron transport (Fig 2B,C, gray and pink clusters) were expected given the hyperpolarization arising from treatment with ribosome inhibitors^19,36^.

The most highly significant GO term of upregulated genes was the phage shock protein (PSP) response. This pathway is involved in sensing and repairing membrane damage^37,38^ with membrane sensors (*pspB, pspC*) and a cytoplasmic aggregation factor (*pspA*), as well as other accessory proteins. All genes from the psp operon increased expression from the earliest time points and were among the largest changes (Fig 2D, Fig S5B). Not all membrane damage sensing pathways^39^ were upregulated during kanamycin treatment, which suggested global membrane stress response was not active (Fig S5B). Previous work has shown aminoglycoside mistranslation induces membrane breakdown^19,32^ and we hypothesized the psp pathway was a mechanism to preserve membrane integrity, thus maintaining the zombie state.

To test this hypothesis, calcium dynamics were measured in knockouts of components of the psp operon treated with gentamicin. The *pspA* knockout cells showed smaller calcium transients (Fig S6A) which ceased more quickly than WT cells (Fig 2E). Other proteins in the psp operon did not have the same pronounced reduction, but Δ*pspF* also stopped transients early (Fig S6B). pspF is the transcription activator for the psp operon, and its knockout prevents the expression of pspA^38^. Knockout of *pspA* also showed an increased rate of PI uptake compared to the WT, consistent with increased membrane damage (Fig 2F). Transmission electron microscopy (TEM) images of WT and Δ*pspA* cells treated with kanamycin for 4 hours revealed large membrane holes in the knockout cells that were much less abundant than WT cells or untreated *pspA* knockouts (Fig 2G, Fig S6C). *Pseudomonas putida*, an unrelated Gram-negative species, does not contain homologs to the psp genes and did not exhibit long-term calcium dynamics after treatment with gentamicin which suggested they were unable to form the zombie state (Fig S7A,B). These cells also showed increased membrane damage after 4-hours treatment with gentamicin when measured by TEM (Fig S7C-E). Thus, we concluded the zombie state is maintained by the psp pathway in *E. coli* which enabled survival for up to 4 days after cell cycle exit, but aminoglycoside-induced zombies are not universal across bacterial species.

### Aminoglycoside-induced zombies increase the antibiotic tolerance of naïve cells

The metabolomic and CFU data showed *E. coli* treated with kanamycin remained metabolically active but were unable to divide, similar to senescence in eukaryotes. One aspect of senescent eukaryotic cells is their secretion profile, which can alter physiology in neighboring cells^40,41^. We hypothesized that zombie *E. coli* could maintain active biochemical synthesis and secretion pathways thereby communicating with neighboring cells. The potential functional consequence would be similar to bacterial charity work, whereby subpopulations of bacteria will produce beneficial signaling molecules at the expense of slower growth^42^.

We sought to test the hypothesis that zombie bacteria produced a signaling molecule that can influence treatment-naïve cells from future antibiotic exposure. Zombie cells were generated with kanamycin treatment for 24 hours, followed by washing 3x with fresh medium to remove the antibiotic (Schematic, Fig S8) and incubating in fresh medium. CFU measurements showed no colonies from zombies but based on the GCaMP6f data (Fig 1C), we expected ∼65% of cells to maintain metabolic activity. The zombies were then mixed with naïve cells that had not been exposed to aminoglycosides. Upon addition of kanamycin, the treatment-naïve cells exposed to zombies were more tolerant than naïve cells without zombies, exhibiting a greater than 30-fold increase in CFUs (Fig 3A). To confirm zombies had not re-entered the cell cycle, carbenicillin-resistant naïve cells were mixed with carbenicillin susceptible zombies. This mixture showed the same CFU counts on both LB and LB-carbenicillin plates (Fig S9A). Conversely, using carbenicillin-resistant zombies with carbenicillin-naïve cells showed no colonies on LB-carbenicillin plates (Fig S9B). Carbenicillin-resistant zombies expressing mScarlet were also absent from re-growth as measured by the absence of pink colonies. Thus, we concluded the zombie cells are not re-initiating the cell cycle upon mixture with naïve cells.

**Figure 3:**
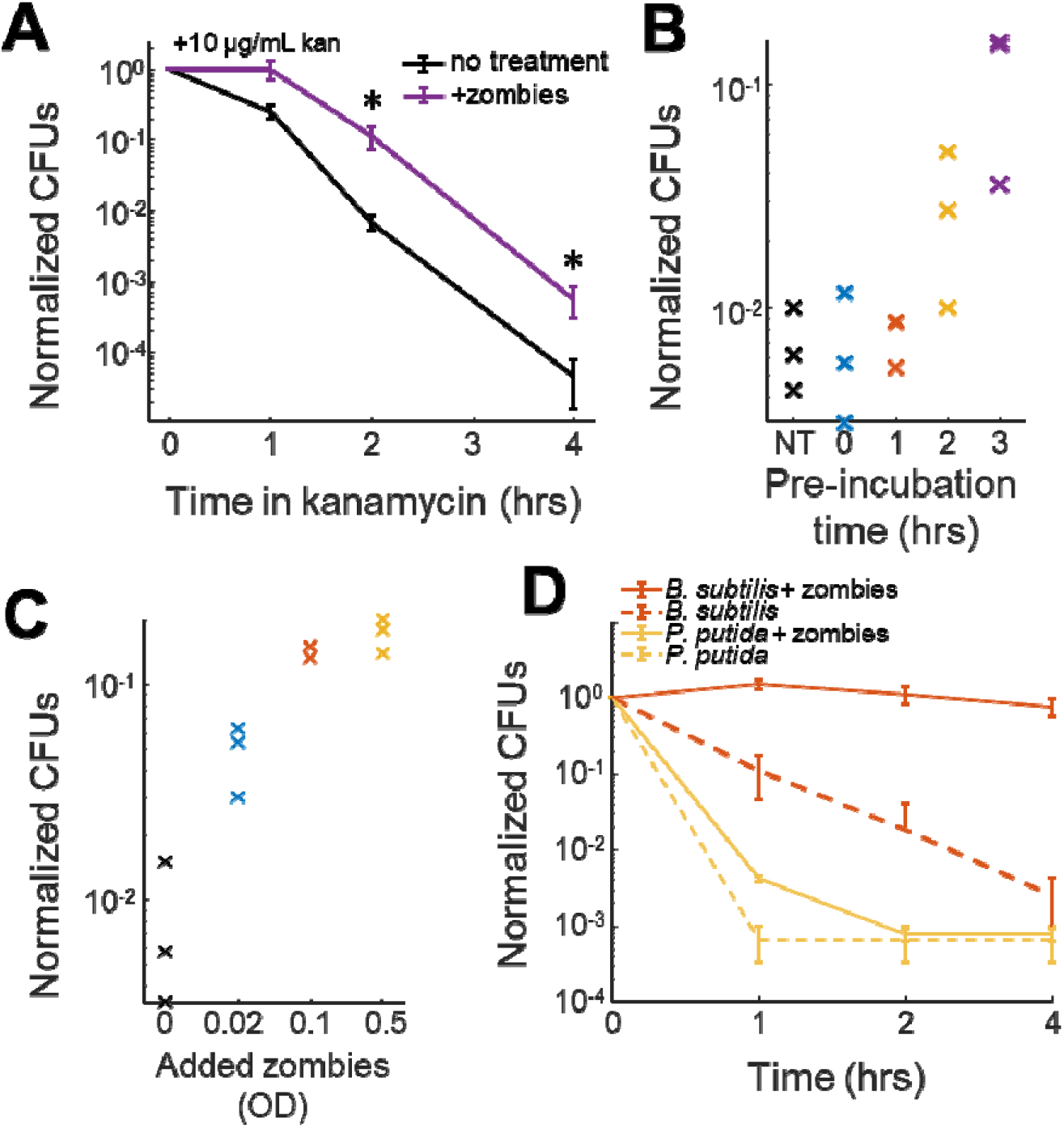
Zombie *E. coli* increase the antibiotic tolerance of naïve bacteria. (A) CFUs of naïve cells treated with 10 μg/mL kanamycin at time t=0 hrs. CFU counts were normalized to the mean of 3 biological replicates at t = 0. Black (no zombies) and purple (with zombies). The error bars are the standard deviation of 3 biological replicates. * denotes p < 0.05 (two-sided t-test). (B) Naïve cell CFUs after 2-hours of treatment with 10 μg/mL kanamycin as a function of the amount of incubation time with zombies. NT i no treatment. Each x is a biological replicate. (C) Naïve CFUs after 2-hours treatment with 10 μg/mL kanamycin as a function of the amount of zombie cells added. The optical density (OD) is the final OD of just the zombie cells. (D) CFUs of naïve *B. subtilis* (red) or *P. putida* (yellow) as a function of tim treated with 10 μg/mL kanamycin. Naïve cells either had no exposure (dashed line) or were incubated with zombies 3 hours before antibiotic addition (solid line). The error bars indicate the standard deviation of 3 biological replicates.

The duration of zombie incubation was critical to the response as less than 2-hours of incubation with zombies did not increase the survival of naïve cells (Fig 3B). At 2 hours or more, increased tolerance was observed. The protective effect of the zombies was dose-dependent (Fig 3C), and heat killed zombies did not protect naïve cells (Fig S10A-C). The heat killed zombies showed the response was not due to residual sub-lethal kanamycin. Naïve cells treated with zombies exhibited an identical growth rate compared to untreated cells in the absence of kanamycin (Fig S10D,E). Thus, we concluded the continued metabolic activity of zombies included secreted factors that can alter antibiotic tolerance.

Many secreted signaling molecules used by bacteria are not species specific^42–44^. Given the potential for interspecies communication, we tested if zombie *E. coli* could protect unrelated bacterial species. Zombie *E. coli* were made as before and added to naïve cultures of *P. putida* or *Bacillus subtilis*. Zombie *E. coli* protected both naïve *P. putida* and *B. subtilis* (Fig 3D). The protection in *B. subtilis* was striking with almost no loss of viability when pre-cultured with zombies, corresponding to an ∼250x increase in CFUs. Combined, our data showed that zombie *E. coli* can communicate with naïve cells and ameliorate their susceptibility to antibiotics.

### Aminoglycoside-induced zombies secrete indole to affect naïve cells

Given that zombies protected naïve cells from antibiotic exposure, we sought to identify the molecular communication used by zombie cells. We first asked if the supernatant from zombie cells was modified. Untargeted metabolomics of the zombie supernatant collected over 96 hours indeed showed a dynamic profile indicating zombies modified their external environment (Figs 4A, S11A, File S5). We then tested if a chemical factor in the supernatant could protect *E. coli*. Similar to directly adding the zombies, we found that the supernatant could protect naïve *E. coli* (Fig 4B) which suggested a diffusible factor.

**Figure 4:**
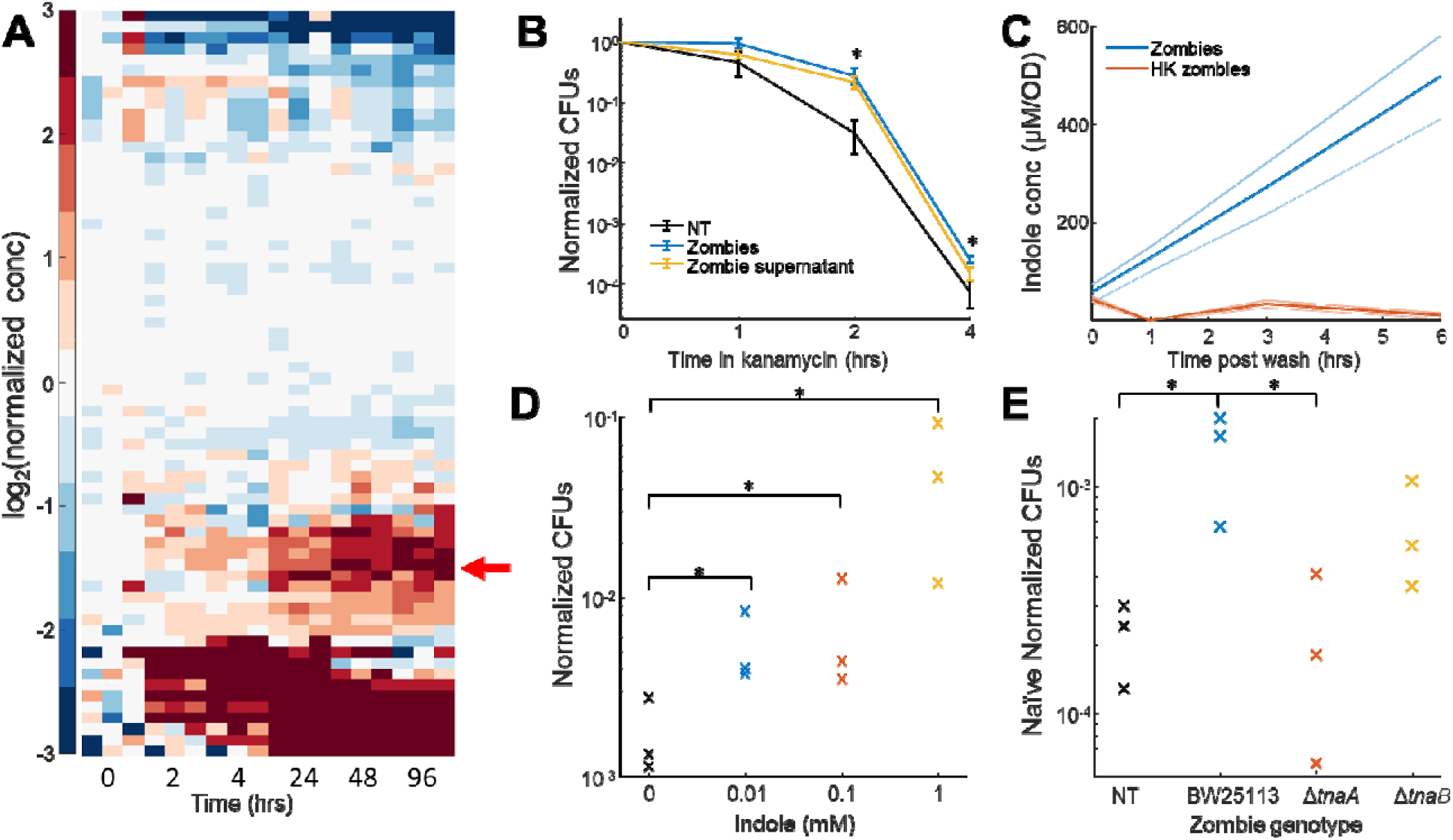
Zombie cells secrete indole to increase naïve cell antibiotic tolerance. (A) Untargeted metabolomic results of the supernatant of *E. coli* when treated with 30 μg/mL kanamycin for the indicated time. Th data was clustered by hierarchical clustering. Red arrow indicates 5-hydroxyindoleacetate (Fig S10B). (B) CFUs measured of naïve cells with no exposure (black), with zombies (blue) or with zombie supernatant (yellow). * denotes p-value < 0.05 (two-sided t-test) when comparing no-treatment to zombie supernatant. (C) Indole production of zombies (blue) compared to heat-killed (HK) zombies (red). Error bars are equal to standard deviation between 3 biological replicates. (D) CFUs of naïve cell treated with 10 μg/mL kanamycin for 2-hours as a function of exogenously added indole. Each x is biological replicate. (E) CFUs of naïve cells treated with 10 μg/mL kanamycin for 2-hours. Naïve cell had no treatment (black), WT zombies (blue), Δ*tnaA* zombies (red), or Δ*tnaB* zombies (yellow). Each x is a biological replicate. In all panels, * denotes p-value < 0.05 (two-sided t-test)

To identify potential signaling chemicals, we examined the supernatant metabolome and noticed increased concentrations of metabolic products in the indole production pathway (Fig 4A red arrow, Fig S11B) which suggested that indole concentration increased with kanamycin treatment, in line with earlier reports^45^. Indole has previously been shown to increase CFUs upon treatment with antibiotics^42,43,46^, so we hypothesized zombie cells could secrete indole as a chemical messenger to influence neighboring naïve cells. We tested if zombie *E. coli* increased medium indole by creating zombies with 24-hour kanamycin treatment, followed by a wash into fresh medium. After treatment, CFU counts were below our detection limit (Fig S12A). Indole concentration, as measured by Kovac’s reagent (Fig S12B,C), increased in the fresh PMM reaching a concentration of 520 μM/OD after 6 hours (Fig 4C). Direct supplementation of indole in the supernatant showed a concentration dependent increase in the number of CFUs upon treatment with kanamycin (Fig 4D). Zombies generated from knockouts of *tnaA* (indole production) and *tnaB* (indole export) showed a reduction in the number of naïve CFUs as compared to WT zombies, consistent with the protective role of indole (Fig 4E). Thus, we concluded that zombies are no longer able to undergo cell division, but continue to secrete indole which can alter antibiotic susceptibility in neighboring naïve cells.

## 4. Discussion

In this work, we showed that *E. coli* that would traditionally be described as dead are, by many definitions, alive and can influence antibiotic susceptibility in neighboring cells belonging to divergent branches of the evolutionary tree. Bacterial survival has long been measured by their ability to form colonies, but this invaluable technique is blind to changes in cell physiology that occur in the absence of cell division. Other types of non-dividing cells have been documented including persisters, VBNC, and minicells^27,33^. Herein we documented a new senescent-like state in *E. coli* treated with aminoglycosides. This state is characterized by permanent cell cycle exit, maintenance of membrane integrity via phage shock response, continued glucose metabolism, and a senescence-associate secretome. Akin to the senescence-associated secretory phenotype in mammals, the zombie secretions were able to chemically communicate with naïve cells via indole and thereby increase the naïve cell’s antibiotic tolerance. Overall, our work indicates that cell death measured by CFUs does not capture the single-cell physiology of antibiotic response; zombies must be accounted for when measuring population-level antibiotic efficacy.

One outstanding question is which mechanisms control *E. coli* entry into the zombie state. Minicells, which exhibit similar phenotypes, arise because the lack of a chromosome necessarily prevents cell growth and division. Zombies retain their chromosomes, but aminoglycosides are known to induce DNA damage^47^, suggesting that irrecoverable DNA damage may trigger the zombie state. This hypothesis would also suggest that other antibiotics classes such as fluoroquinolones or folic acid inhibitors could also induce *E. coli* or other bacterial zombies. We believe that identifying the mechanisms underlying the transition into zombies will reveal other environmental perturbations that induce this state.

Zombies secrete paracrine signals that can alter the antibiotic tolerance profiles of naïve cells from diverse bacterial species. Therefore, they may tune the consortium composition upon a second dose of aminoglycoside, as those more responsive to indole, such as *B. subtilis*, will have an advantage compared to indole non-responsive species. Another possibility arising from zombie signaling is the potential for direct communication with host cells via secretion of indole, LPS shedding, or other small molecules^48,49^. Finally, the emergence of altruistic phenotypes in particular bacterial orders, which cannot be genetically selected, poses an interesting case study for investigating how such phenotypes arise.

## Acknowledgments

This study was funded by the Searle Scholars Program and NIH New Innovator award (1DP2GM123458) to JMK. CTM is funded from T32AG000279. GNB was funded through the T32 training grant (T32GM065103) and HHMI Gilliam Fellowship for Advanced Study. The Anschutz metabolomics core is supported by the Cancer Center Support Grant P30CA046934. Electron microscopy was done at the Boulder EM Services Core Facility in the MCDB Department, with the technical assistance of facility staff. Flow cytometer was used in the Biochemistry Shared Instrument Pool at CU Boulder and was acquired with an instrumentation grant (NIH S10OD021601). We thank Megan Jewell for help preparing cells for electron microscopy and for expressing GCaMP6f in *P. Putida*. We thank Jacob Fenster for protocols for growing *P. putida*. The authors declare no conflict of interest. The funders had no role in the study design, data collection, data analyses, writing, or decision to publish.

## Materials and Methods

### a. Strains and plasmids

Strains used in these experiments were *E. coli* BW25113 (from Yale Cole Genetic Stock Center), *P. putida* KT2440 (provided by the Cameron Lab), and *B. subtilis* W168 (provided by the Garner lab). Knockouts were acquired from the Keio collection (Dharmacon, OEC4988). The GCaMP6f-mScarlet and GCaMP6f alone plasmids were produced previously^15^ and used under a constitutive promoter. The *P. putida* expression plasmid was constructed using a pBBR1 broad host origin plasmid that was cut with restriction enzymes and the 118 constitutive promoter from *E. coli* was used to drive expression. The GCaMP6f-mScarlet ORF was cloned behind the 118 promoter, and the plasmid was transformed into *P. putida* using electroporation. Cells harboring the pBBR1 plasmid were grown in 50 μg/mL kanamycin to maintain expression.

### b. Cell growth

*E. coli* and *B. subtilis* were plated from a glycerol stock onto LB-agar plates with an appropriate resistance (no antibiotic, 100 μg/mL carbenicillin, 50 μg/mL kanamycin) and grown overnight in a 37 °C standing incubator. From these plates, individual colonies were picked and grown in 4 mL LB with antibiotic and grown overnight shaking at 37 °C, reaching stationary phase full saturation by morning. From the overnight culture, cells were diluted 1:100 into PMM (1x M9 salts, 0.2% glucose, 0.2 mM MgSO_4_, 10 μM CaCl_2_, 1x MEM amino acids, pH 7.5). These cells were placed into a shaking incubator at 30 °C.

*P. putida* were grown similar to *E. coli*, except the overnight growth was performed at 30 °C.

### c. Imaging experiments

Imaging experiments were performed similar to previous work^15,19^. First, agarose pads were prepared using a 3D printed mold with dimensions that fit into a 96 well glass bottom dish. The mold was designed so that the final volume of agarose in each well is 200 μL. The agarose pad was made with PMM and 1% agarose. Cells from the diluted stock in PMM were grown for 2 hours at 30 °C, and 2 μL of this cell suspension was placed onto the agarose pad. The culture was left to absorb into the pad for 15 minutes ensuring no excess liquid.

The cells on the agarose pad were then pressed into a 96 well glass bottom dish (Brooks, Matrical). One field of view (FOV) was selected for each well to be imaged before the addition of any compounds. Compounds were added by placing a 5 μL drop on top of each agarose pad, where each drop was at 40x the final concentration. Measurements show compounds reach the sample within 5 minutes and equilibrate to a final concentration within 30 minutes^20^.

Imaging 96-well glass bottom plates took place using a Nikon Ti2 inverted microscope running the Slidebook software (3i). The temperature of the sample was held at 30 °C using a microscope top incubation chamber (Okolabs). Fluorescent excitation was achieved with a solid state laser illumination. A 40x, NA 0.95 air objective was used to both illuminate and image the cells onto 2-Prime95B sCMOS cameras (Photometrics) using a Cairn image splitter to image two colors. Illumination was achieved by sequential excitation with 488 or 561 nm laser for a 200 ms exposure. Measured light intensities at the sample were 180 mW/cm^2^ (488 nm) and 450 mW/cm^2^ (561 nm). Typical sampling rates were one frame per minute, unless noted in the text. At a frame rate of 1 per minute, we could typically image 32 wells at most. The perfect focus was used to maintain the focal plane over 4 days of imaging, though some drift was unavoidable in our setup.

Samples using propidium iodide (PI) were performed by dissolving 3 μg/mL PI into the agarose pad before pouring into the 3D printed mold. These measurements were performed using cells expressing GCaMP6f alone.

Heat killing of bacterial cells was performed using a heat block set to 65 °C. A volume of < 500 μL was placed into a 1.5 mL centrifuge tube and set in the heat block for exactly 10 minutes. The cells were then removed and either washed, or placed directly onto agarose pads for imaging experiments.

### d. Image processing

Image processing of GCaMP6f movies was performed similar to previous work^19–21^. All analysis was carried out using custom scripts in Matlab (Mathworks, R2020b). To estimate an uneven illumination background, we loaded 10 frames from each FOV and calculated the mean pixel-wise intensity. This was blurred using a Gaussian kernel of 55 pixels and all FOVs were averaged together. This smoothly varying image was used as our estimate for the illumination pattern.

To process an individual FOV, the first 10 frames were loaded into memory and the average image was used as the basis for an XY registration. Segmentation was performed on the average of the first 10 frames using a Hessian to detect lines.

Then, each frame was loaded to memory one at a time and went through the following processing: 1) Divide by the illumination pattern and multiply by the mean of the illumination, 2) register the current image to the basis image using a FFT alignment, 3) subtract the background using a structured element to identify local background, 4) remove any pixels that have drifted out of the imaging FOV, and 5) extract the fluorescence intensity from every cell that remains entirely in the imaging plane. Due to the long imaging times (96 hours) there were movies that drifted entirely out of the field of view. These movies were discarded.

To identify the cessation of calcium dynamics, all the traces from a given experiment were randomized and displayed to the user. We manually chose the time at which there was no visible activity afterwards. We note our approach could underestimate time to cessation for some cells that re-established dynamics at times > 96 hours. The timings were then re-sorted back into each condition, and the estimated cumulative distribution function is plotted.

### e. CFUs

CFUs were measured using 10x dilution spot plate assay. Briefly, from each culture, ∼200 μL was taken and put into a 96 well plate. The cultures were diluted by taking 20 μL and diluting into 180 μL each row. A 12 channel electronic pipet was set to the pipet and mix function, and new tips were used after each 10x dilution step. From this dilution series, 3 μL was plated onto an agar-LB plate. Colonies were counted by identifying the series with at least 8 colonies, and multiplying by the appropriate 10x dilution constant.

### f. Flow cytometry

Median TMRM values were obtained using flow cytometry. WT *E. coli* from an overnight culture were diluted 1:100 into PMM and incubated at 30 °C for 2h. After incubation, kanamycin sulfate in dIH_2_O (5 mg/mL) was added to each sample for a final concentration of 30 µg/mL kanamycin. Samples were prepared 96h, 48h, 24h, and day-of measurement, then analyzed during the same flow cytometry session. One hour before flow measurement, CCCP was added to negative controls for a final concentration of 50 µM. Thirty minutes before measurement, tetramethylrhodamine methyl ester (TMRM, 20 µM in DMSO) was added to every sample for a final concentration of 200 nM. Flow cytometry measurements were performed on a BD FACSCelesta flow cytometer.

Cells were quantified for their TMRM incorporation by counting 100,000 events per condition using a BDFACSCellesta Flow Cytometer with the following voltage settings: FSC at 700, SSC at 350, with 561 nm laser D585/15 at 500, C610/20 at 500 and B670/30 at 481. Emission for each event was collected at the 585/15 nm wavelengths.

### g. Metabolomics

To generate cells and supernatant for metabolomic experiments, cells were grown overnight in LB as above, and diluted into 40 mL PMM at a concentration of 1:100. A total 3 biological replicates were identically diluted into 6x 40 mL tubes and placed into a shaking incubator at 30 °C. After 2 hours, 30 □g/mL kanamycin was added to each tube. At time points of 0, 2, 4, 24, 48, and 96 hours, cells and supernatant were collected.

To collect the supernatant and cells, the tubes were removed from the shaking incubator at the appropriate time.

Supernatant - first 400 μL was taken off and placed into a centrifuge tube. This sample was spun down at 19000x g for 2 minutes. A 200 μL aliquot of the supernatant was then flash frozen in a fresh tube in a mixture of ethanol and dry ice for 1 minute, and then placed into a -80 °C freezer.

Cells – The centrifuge was first cooled to 4 °C. After removing the 40 mL tube from the incubator and the supernatant sample was removed, the 40 mL tube was placed into the dry ice/ethanol bath for exactly 65 seconds. This amount of time was enough to rapidly cool the sample but not form ice crystals. The cooled sample as then spun down at 3000x g for 3 minutes to form a cell pellet. The supernatant was decanted and the cell pellet was washed in ice cold PBS. The cells were again spun at 3000x g for 3 minutes and the supernatant was decanted. The cell pellet was then flash frozen for 1 minute in the dry ice/ethanol bath, and the samples were placed into a -80 °C freezer.

Mass spectrometry metabolomics was performed by the CU-Anschutz Mass Spectrometery Metabolomics Shared Resource Facility. An ultra-high performance liquid chromatography-mass specrometery run of polar metabolite profiles was performed on both supernatant and cell pellet samples.

Analysis of metabolomic data was performed in Matlab (R2020b). Each metabolite was normalized to the mean of the t=0 hrs timepoint. The normalized data was then clustered only along the concentration dimensions (not time) with a correlation distance measure using the *clustergram* function in Matlab. The principal component analysis was performed on the normalized data set.

### h. Glucose detection

To collect cells for glucose measurements, first cells were grown overnight in LB and diluted 1:100 in PMM as above. These cells were incubated at 30 °C for 2 hours and 30 μg/mL kanamycin was added. The cells were then incubated for 24 or 48 hours at 30 °C to ensure zero CFUs. These cells were then washed 3x in fresh PMM. At this time point, and at 4, 24, and 48 hours after washing the supernatant was collected by spinning down 200 μL, and was flash frozen in a dry ice/ethanol bath.

To measure glucose consumption in the medium, an Amplex Red Glucose detection kit was used according to manufacturers’ directions (LifeTech, A22189). The Amplex Red reaction mixture was prepared directly before use. The supernatant from treated cells was added to each well (50 μL) in a 96 well plate, followed by the addition of 50 μL of the Amplex Red reaction mixture. This mixture was allowed to incubate for 30 minutes at room temperature, followed by fluorescence measurement using the Tecan Spark plate reader.

### i. RNA isolation and sequencing

To generate material for RNAseq, cells were grown in LB and diluted in PMM as above. To isolate RNA, we used a modified version of the RNASnap protocol^50^. Briefly, we used a mixture of formamide, 2-mercapto-ethanol, SDS, and EDTA as the RNAsnap buffer. We used 750 μL with a 750 μL cell culture.

To collect the cell pellet, treated cells were added to a pre-chilled centrifuge tube containing 750 μL cold methanol. These tubes were then spun down in chilled centrifuge for 3 min at 18,000x g. The supernatant was decanted and the cell pellet was then flash frozen in liquid nitrogen or directly moved to cell lysis. To lyse cells, the pellet was resuspended in 100 μL of RNAsnap buffer, incubated at 95 ºC for 7 min, and centrifuged at 18,000x g for 5 minutes. The supernatant was carefully transferred to a fresh tube leaving behind the cell debris.

The RNA was then cleaned up, with an on column DNAse treatment, using a Zymo RNA Clean and Concentrator 25 column using the manufacturer’s protocols. RNA was eluted with 35 μL molecular biology grade water. The RNA was quantified using a nanodrop and bioanalyzer. RIN values > 8 were obtained for all the samples.

RNA libraries were prepared using the Illumina bacteria single stranded total RNA prep. Samples were depleted of rRNA using ribozero according to the manufacturers’ directions. Sequencing was performed on an Illumina NextSeq500 with a total of > 10M reads per sample.

### j. RNAseq analysis

RNAseq analysis was first performed by the bacterial genomics software, Rockhopper^51^. FastQ files were assigned experiments within the software and the genome sequence was set to *E. coli* BW25113. The Rockhopper software returned RNA counts for each gene which were fed into the differential expression and clustering algorithms (File S2).

To identify gene clusters in time-series data, accelerated Bayesian Hierarchical Clustering (BHC) was employed^34,35^ using the available R package. Initially, genes were filtered by expression using the *filterByExpr* command in the R package *edgeR*. Subsequently, differentially expressed genes (DEGs) were computed between groups defined by the 6 time-points. Pair-wise algorithms for identifying DEGs (e.g. *edgeR*) were found to outperform algorithms specific for time course clustering when considering time-series of fewer than 8 time-points^52^. Genes with an adjusted p-val<0.001 were included in subsequent analysis resulting in 1746 DEGs (File S3). From this subset, the gene clusters were computed using the *bhc* package in R. Cluster number was selected based on visual inspection. Gene Ontology (GO) analysis^53^ was performed using the PANTHER tool^54^. Due to the small size of the *E. coli* genome, no multiple hypothesis correction was applied to the resulting p-values. See File S2 for genes in each cluster and File S3 for significant ontologies for each cluster. The number of genes in each cluster is annotated in File S2. Dataset accession has been created in GEO (GSE184377).

### k. Transmission electron microscopy

Cells to be imaged with TEM were grown in LB overnight and then diluted into PMM at 1:100 as above. These cells were grown for 2 hours at 30 ºC. At this point, the untreated cells (BW25113, Δ*pspA*, and *P. putida*) were removed, and 30 μg/mL gentamicin was added. These cells were incubated for 4 hours at 30 °C, and then the treated cells were removed.

Samples were pelleted to a loose pellet and supernatant discarded. A few microliters of pelleted sample (per shot) were high pressure frozen using a Wohlwend Compact 02 high pressure freezer (Technotrade International, Manchester, NH) as described previously^55^. Frozen specimens were then freeze-substituted in anhydrous acetone containing 2% osmium tetroxide and 0.2% uranyl acetate and embedded in Epon/Araldite resin. Serial thin sections (80 nm) were cut using a Leica UCT ultramicrotome. The serial sections were collected on Formvar-coated copper slot grids and post-stained with 2% aqueous uranyl acetate followed by Reynold’s lead citrate. The samples were imaged in a Tecnai T12 Spirit TEM, operating at 100 kV using an AMT CCD camera. Greater than 80 cells per condition were counted for the analysis.

#### 1. Zombie protection assays

##### Zombie preparation

*E. coli* BW25113 expressing mScarlet118 were grown overnight in LB as above, and diluted 1:100 in 8 mL of fresh PMM. These cells are then grown in a 30 ºC shaking incubator for 3 hours. Kanamycin (or gentamicin when using the Keio collection) was added to a final concentration of 30 □g/mL. The cells were left incubating at 30 °C for 18 hours.

The next day, cells were spun down at 2250x g for 10 minutes and washed in fresh PMM 3x. If cells were heat killed, they were placed into the hot plate at 65 °C for 10 minutes after the second wash. Zombies were re-suspended in 1 mL fresh PMM to be used in protection assays. The OD of the final suspension was typically around 0.8 OD/600 nm. These cells in the fresh PMM were then placed into the 30 °C shaking incubator for 1 hour.

##### Protection assays

Naïve wild-type cells are prepared in the same manner as described in the CFU assay section above. The primary difference is that 3h prior to antibiotic addition, re-suspended zombies or filtered supernatant is added to the naïve cells. The CFU assays were then performed via the 10x dilution series as described earlier. Unless otherwise noted, CFUs were counted at t = 0, 1, 2, and 4 hours after addition of kanamycin. Spot plates are checked for red colonies (evidence of mScarlet presence and zombie contamination) before colonies are counted. Spots were also plated onto a carbenicillin plate to ensure that any re-growing colonies do not have the GCaMP6f plasmid indicative of a zombie.

### m. Indole quantification assays

Indole was quantified in liquid cultures via the Kovacs assay. Briefly, 150 μL of Kovacs reagent (4-dimethylaminobenzaldehyde and hydrochloric acid solution in n-butanol; Sigma Aldrich) was placed into a 96 well clear bottom plate for each sample to be measured. 150 μL of cell suspension was removed from the shaker and spun down at 19000x g for 2 minutes to pellet any cells. From the supernatant, we removed 100uL and mixed with the Kovac’s reagent in the 96 well plate, pipetting up and down 5x. The samples were then incubated at room temperature and the absorbance spectrum was measured for each well using the Tecan spark plate reader. The absorbance was measured from 450 to 700 nm using a bandwidth of 2.5 nm.

To create a standard curve, each absorbance spectrum was first subtracted by a blank well (fresh PMM, no indole). The corrected sample was then truncated between 491 and 590 nm was integrated in Matlab using the *trapz* function. The three biological replicates were then fit to a line with an R^2^ of 0.983. The resultant linear fit was then used for the measured samples to quantify the amount of indole in the supernatant experiments.

