## Supplemental Materials for "Aminoglycosides induce a bacterial senescent state that increases antibiotic tolerance in treatment-naïve cells"

**Supplementary materials:**

Supplementary movie legends – 1

Description of supplementary files – 1-5

Supplementary figures – 1-12

**Supplementary movie legends:**

Supplementary movie 1: Movie of *E. coli* expressing GCaMP6f exposed to 30 μg/mL kanamycin at time t = 0. The time in the top left is given in DD:HH:MM. The movie was acquired at a frame rate of 1 frame per 2 minutes. The movie began to lose focus around t = 70 hours, but cells could still be seen throughout.

**Supplementary file description:**

Supplementary file 1: Results of untargeted metabolomics of *E. coli* cells treated with 30 μg/mL kanamycin at 0, 2, 4, 24, 48, and 96 hours post treatment. Each metabolite is presented from 3 biological replicates.

Supplementary file 2: RNAseq counts of *E. coli* treated with 30 μg/mL kanamycin at 0, 30, 90, 240, 360, and 1440 minutes post treatment. Each gene is shown with three biological replicates. Counts are given in CPM.

Supplementary file 3: List of differentially expressed genes upon kanamycin treatment. See methods for calculation of DEGs across time.

Supplementary file 4: GO terms from each gene expression cluster.

Supplementary file 5: Results of untargeted metabolomics of *E. coli* supernatant treated with 30 μg/mL kanamycin at 0, 2, 4, 24, 48, and 96 hours post treatment. Each metabolite is measured for 3 biological replicates.

**Supplementary figures:**

Figure S1:

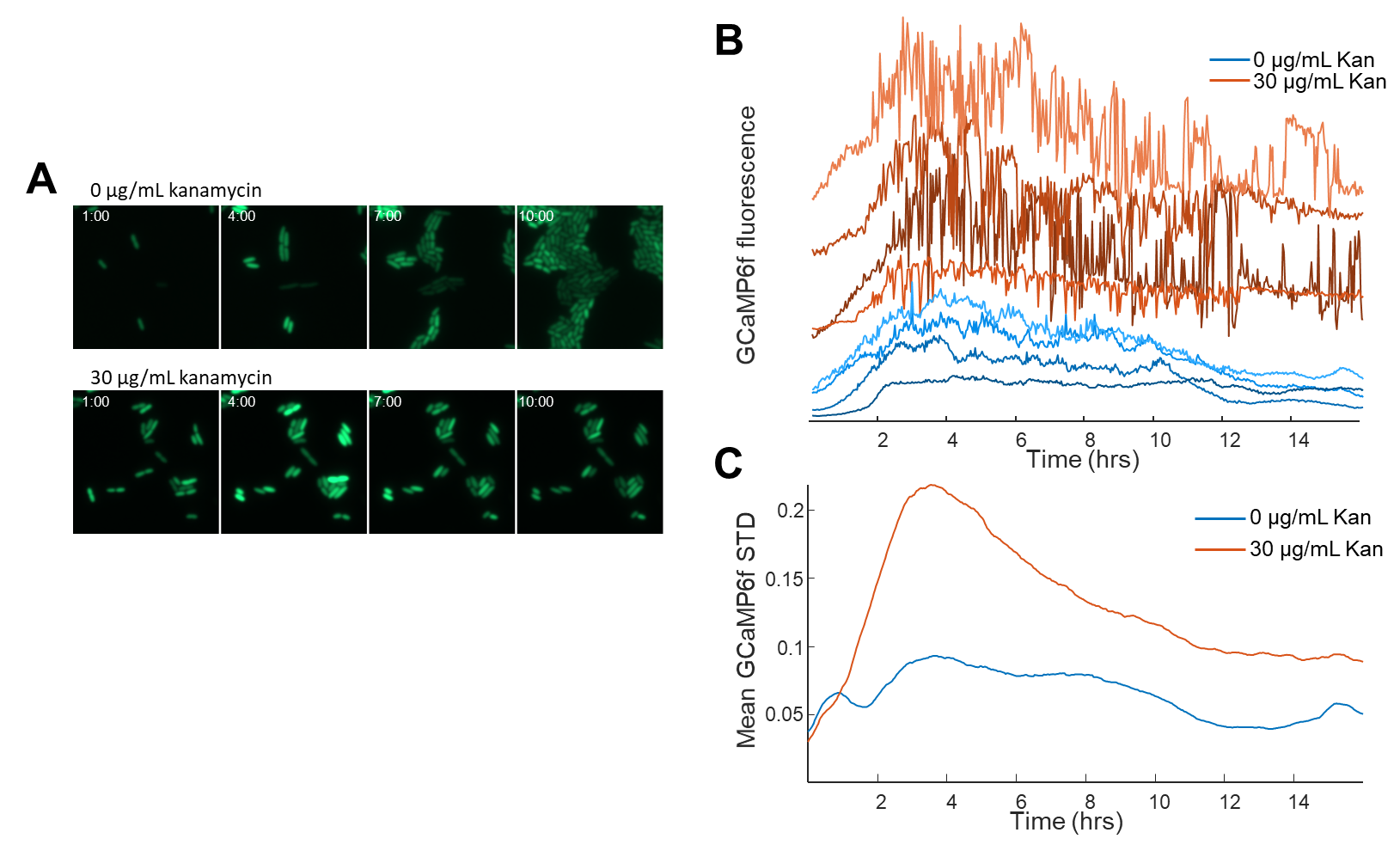

Kanamycin induces cessation of growth and dysregulated calcium for extended times. (A) Strip-chart images of *E. coli* imaged via GCaMP6f fluorescence. The time shown is HH:MM. The cells without antibiotic continue to grow, whereas treated cells are growth arrested after 4 hours. (B) Example traces from single cells either without (blue) or with (red) 30 μg/mL kanamycin. Kanamycin was added at time t=0. The calcium transients in the red traces are indicative of continued metabolism throughout the 16-hour movie. (C) Average moving standard deviation (STD) of all cells with 0 (blue) or 30 (red) μg/mL kanamycin.

Figure S2:

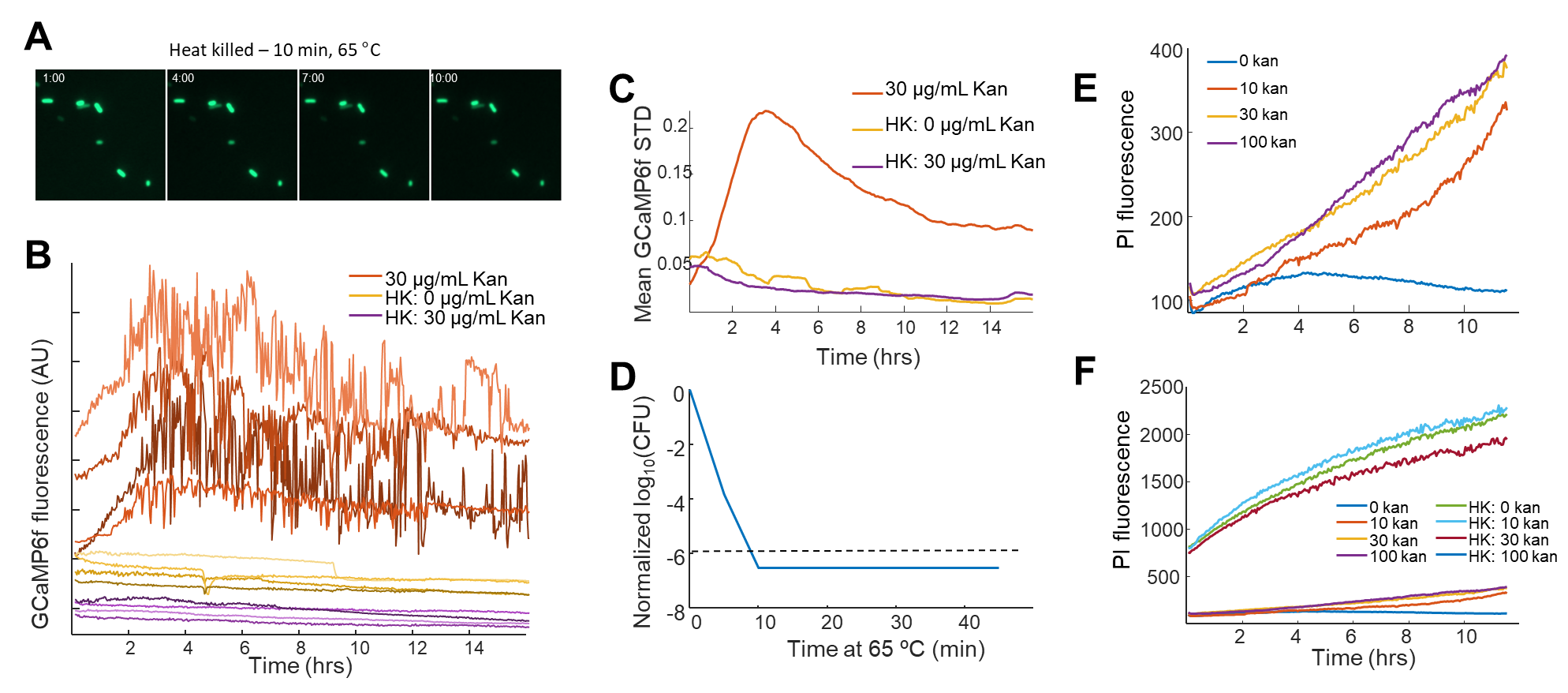

*E. coli* treated with aminoglycosides remain membrane integrity. (A) Image of heat killed cells expressing GCaMP6f. The time is given in HH:MM. The cells maintain fluorescence indicating membrane integrity enough to retain proteins in the cytoplasm. (B) Fluorescence traces from individual cells either treated with kanamycin (red), heat killed (yellow), or heat killed and kanamycin (purple). Heat killed cells do not show any evidence of calcium transients when compared to live cells treated with aminoglycosides. (C) Mean moving standard deviation of the conditions from B. (D) CFUs of *E. coli* as a function of time when heat killed. After 10 minutes at 65 °C, the number of viable cells drops below our detection limit (333 cells/mL). (E) Mean propidium iodide (PI) uptake for *E. coli* treated with 0, 10, 30, or 100 μg/mL kanamycin. PI uptake results from a destabilized membrane that allows the charged fluorophore to enter the cell and bind to DNA. (F) The same data as in (E), except also plotted are heat killed cells. The heat killed cells all take up > 12x propidium iodide as compared to aminoglycoside treated cells. Thus, though aminoglycoside cells do take up some propidium iodide, they remain largely intact compared to heat killed cells.

Figure S3:

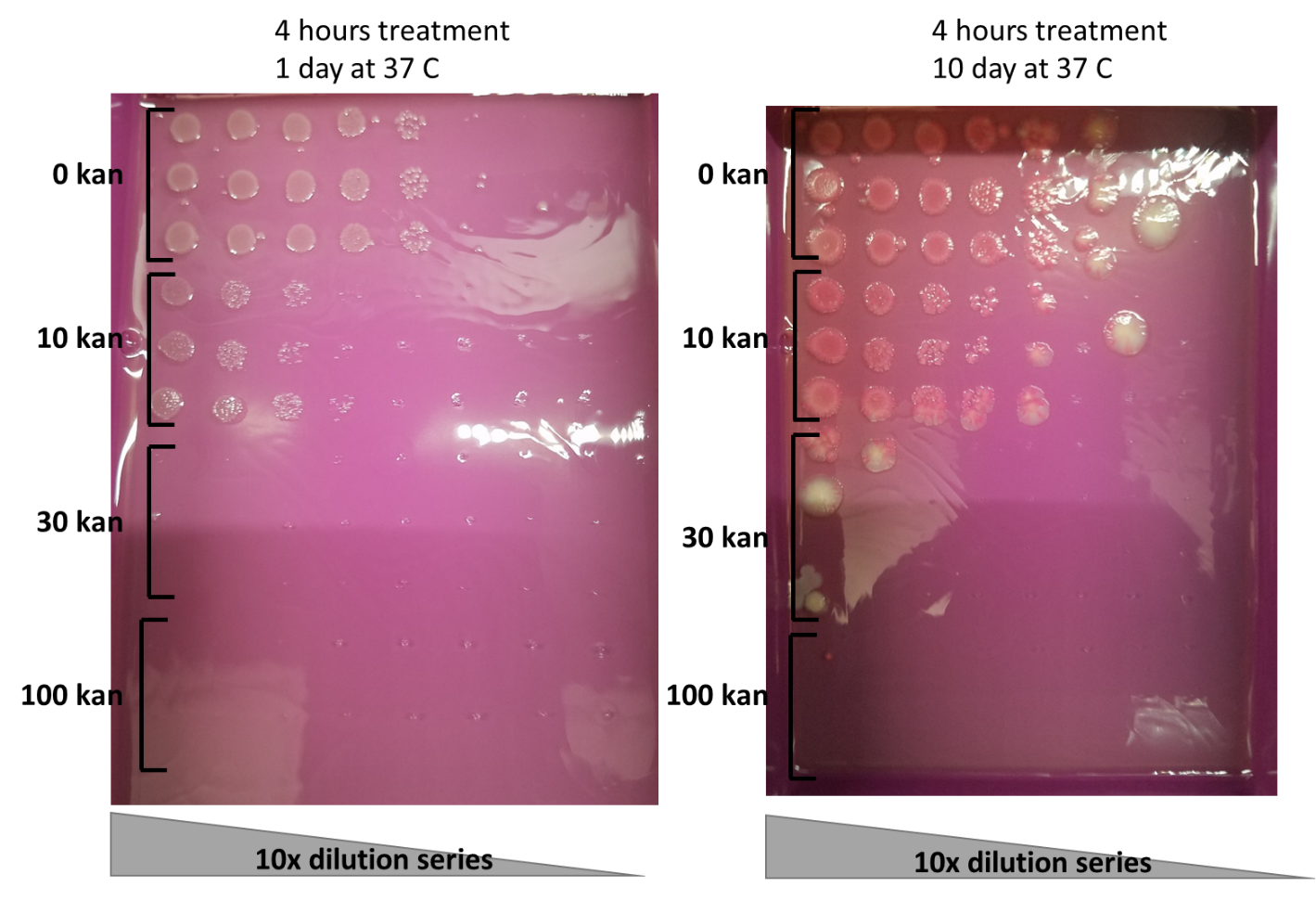

Aminoglycoside treatment is bactericidal, not bacteriostatic. Image of the same CFU spot plate after 4 hours treatment with the indicated amount of kanamycin. 3 biological replicates (rows) per dilution series (columns). Left – spot plate after 1 day at 30 ºC. Right – same spot plate after 10 days at 30 °C. No extra colonies grew after 10 days on the permissive LB+agar plate, indicating that cells are truly unable to divide after 4 hours treatment. If kanamycin was bacteriostatic and aminoglycoside-treated cells could re-enter the cell cycle, we would expect to see new colonies forming with time on the rich medium.

Figure S4:

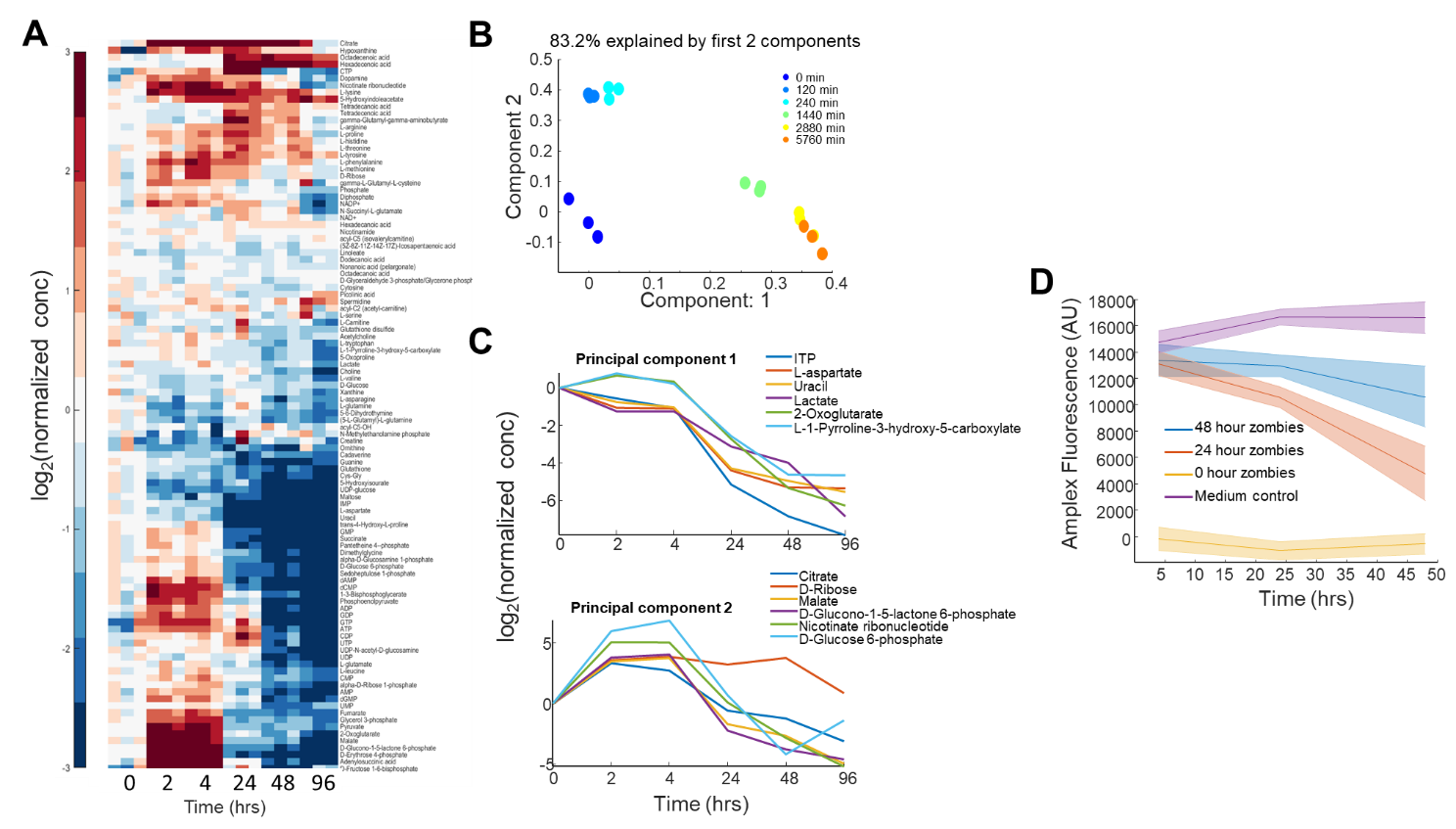

The metabolic profile of cells continues to change over 96-hours of aminoglycoside treatment. (A) Cells continuously treated with 30 μg/mL kanamycin were measured using untargeted metabolomics mass spectrometry. The changes in metabolites over time were clustered using a hierarchical clustering algorithm on just the rows. The heat map shows change in metabolite profile across 96 hours. Each metabolite was normalized to the mean value of 3 biological replicates at time t = 0 hours (before treatment). (B) PCA decomposition of the metabolomic profile at each time point. The color represents the time point showing good clustering and changing profiles with time in aminoglycoside. (C) The metabolites with the largest eigenvalues for the first two principal components as a function of time. Each metabolite is the mean of 3 biological replicates and is normalized to t = 0. (D) Extracellular glucose consumption of zombies as measured by Amplex Red fluorescence. Zombies were created by either 24 hours (red) or 48 hours (blue) of continuous kanamycin treatment before being re-suspended in fresh PMM. Yellow indicates untreated cells, and purple is medium alone. The untreated cells consumed all the available glucose within the first 4 hours before the measurement. The rise in the medium alone (purple line) is indicative of evaporation in the plate over the 48 hour experiment.

Figure S5:

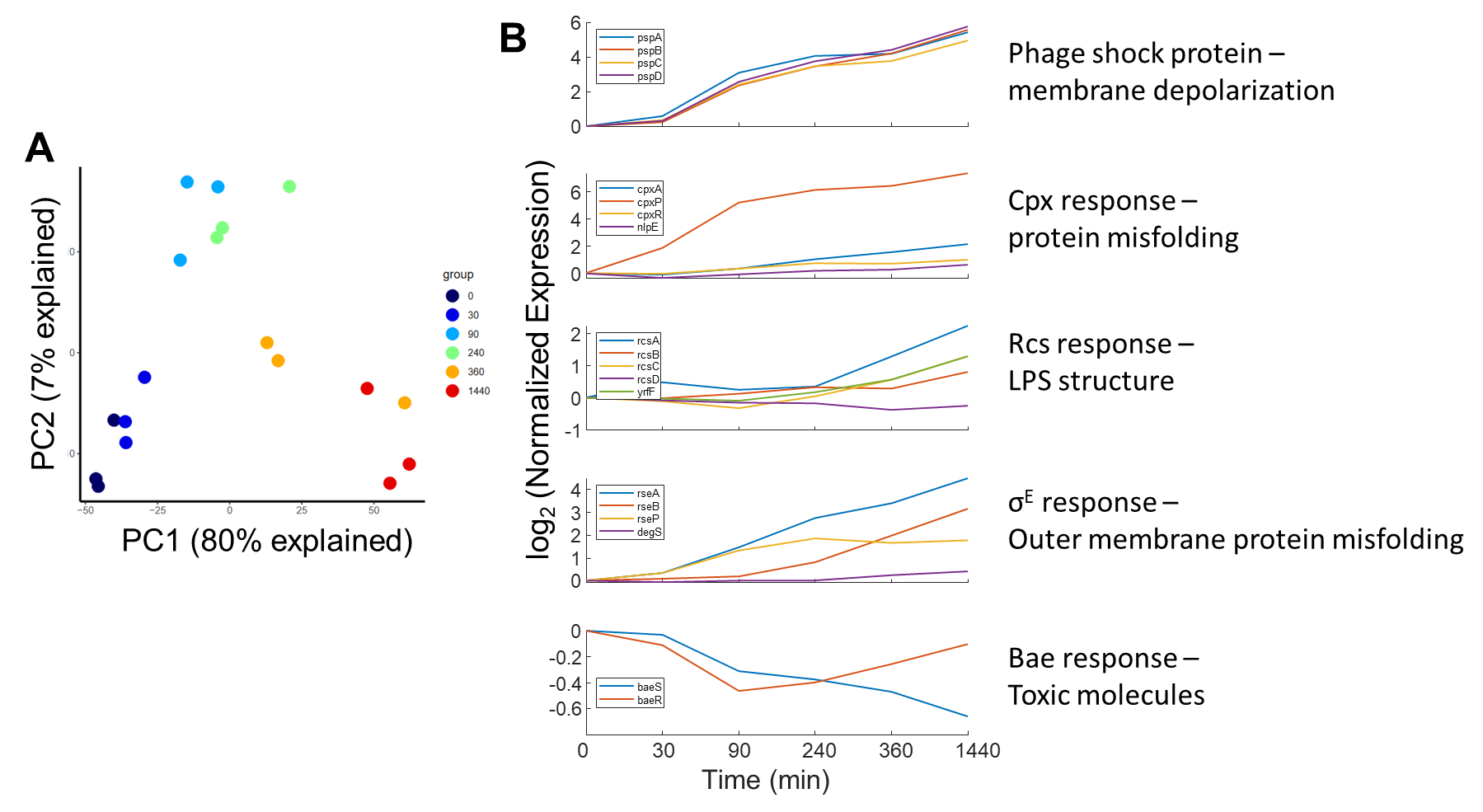

The gene expression profile of zombies changes with time after kanamycin treatment. (A) PCA decomposition of the DEGs (File S3) at each time point. The timepoint replicates (colors) cluster while variability between timepoints shows a two-stage trajectory. First, early global changes occur within the first 240 min (PC 2) followed by orthogonal expression changes for timepoints > 360 min (PC 1). Time is given in minutes. (B) Change in RNA expression of membrane damage sensing pathways as a function of time of aminoglycoside treatment. Only the phage shock protein response (top panel) was universally upregulated over the timecourse of treatment.

Figure S6:

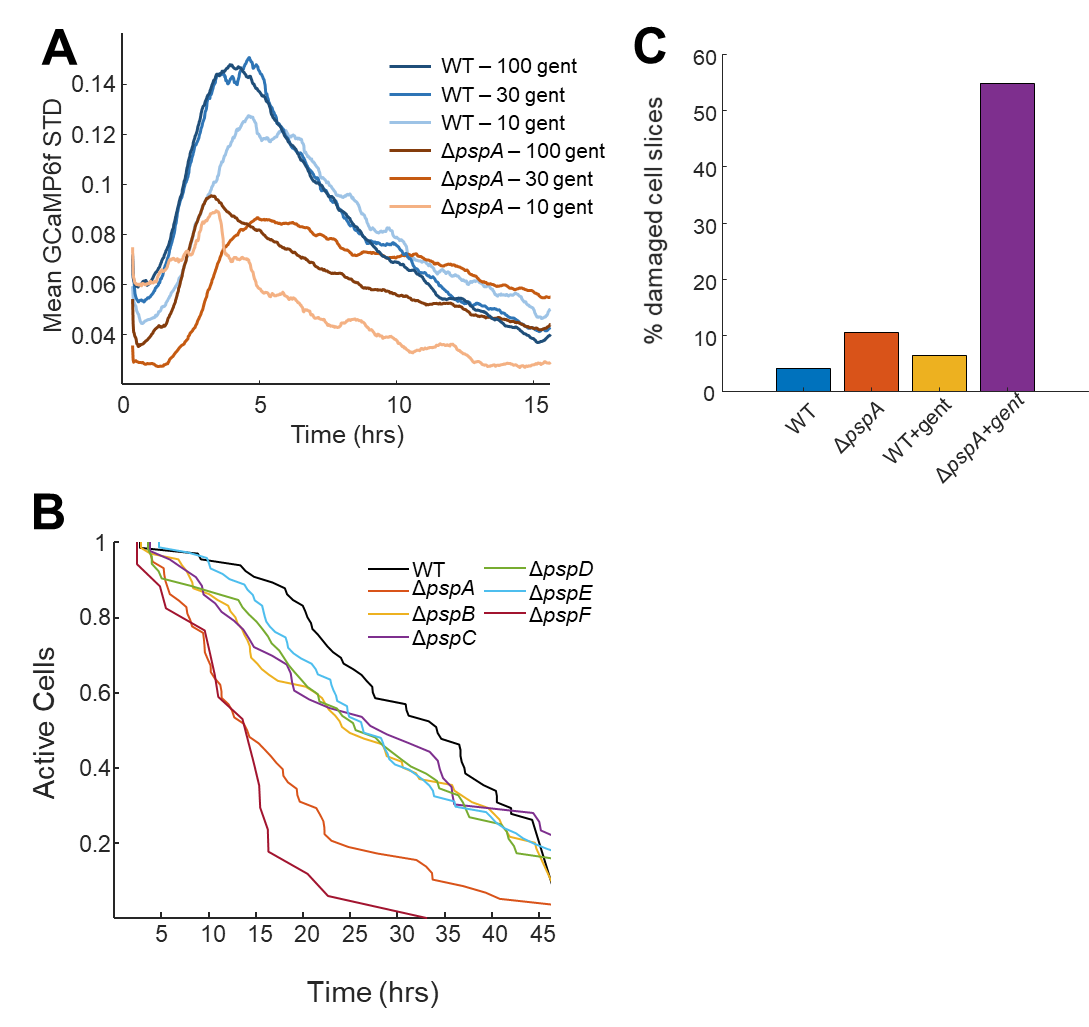

The phage shock protein response maintains membrane integrity upon aminoglycoside treatment. (A) Mean moving standard deviation of GCaMP6f fluorescence from WT (blue) or Δ*pspA* cells treated with gentamicin (legend). Δ*pspA* cells showed smaller calcium transients that decreased more rapidly than WT cells, suggesting they were not able to maintain membrane integrity as long. (B) Kaplan-Meyer curve for knockouts of the psp operon when treated with gentamicin. Cessation of calcium dynamics occurred earlier for Δ*pspA* (orange) and Δ*pspF* (red) as compared to WT (black) and other knockouts of the operon. (C) Fraction of *E. coli* cells with visible damage in an 80 nm slice as measured by TEM. This count likely underestimates the amount of damage as only a thin slice of each cell is visible. The cells were measured at 4 hours post treatment.

Figure S7:

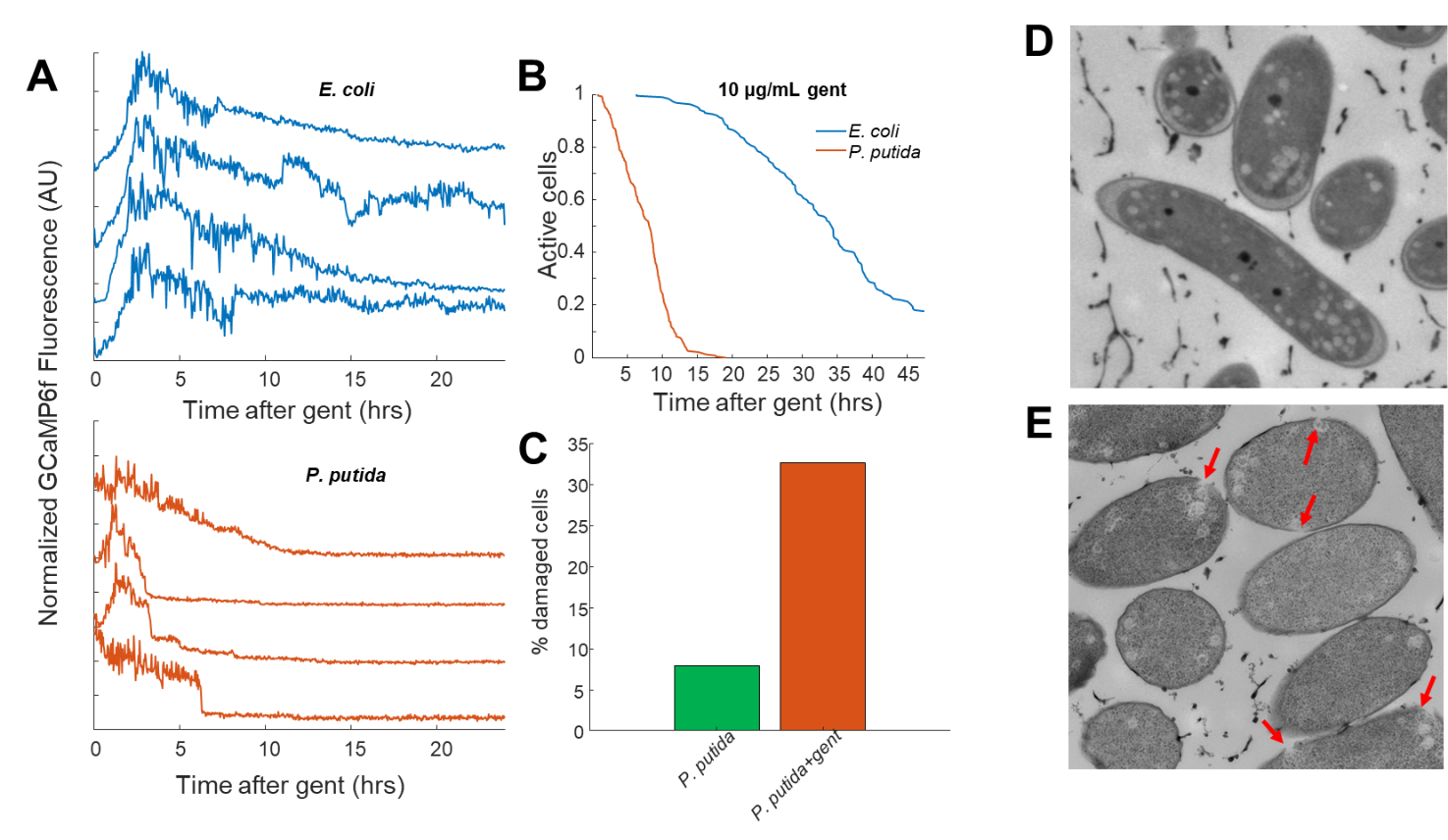

*P. putida* do not form zombies upon treatment with aminoglycosides. (A) Single cell traces of GCaMP6f upon treatment with 10 μg/mL gentamicin at t = 0 hrs. Each trace is an individual cell from *E. coli* (blue) or *P. putida* (red). The *P. putida* cells cease calcium transients earlier than *E. coli*. (B) Kaplan-Meyer curve of activity cessation for *E. coli* (blue, n=186 cells) and *P. putida* (red, n=207 cells). (C) Fraction of *P. putida* cells exhibiting membrane damage before (blue) and after (red) 4 hours treatment with kanamycin. (D,E) TEM micrographs of *P. putida* that were untreated (D) or treated with 10 μg/mL kanamycin (E) for 4 hours. The red arrows indicate sites of membrane damage in the treated condition that are consistent with *P. putida* lacking homologs to *pspA*.

Figure S8:

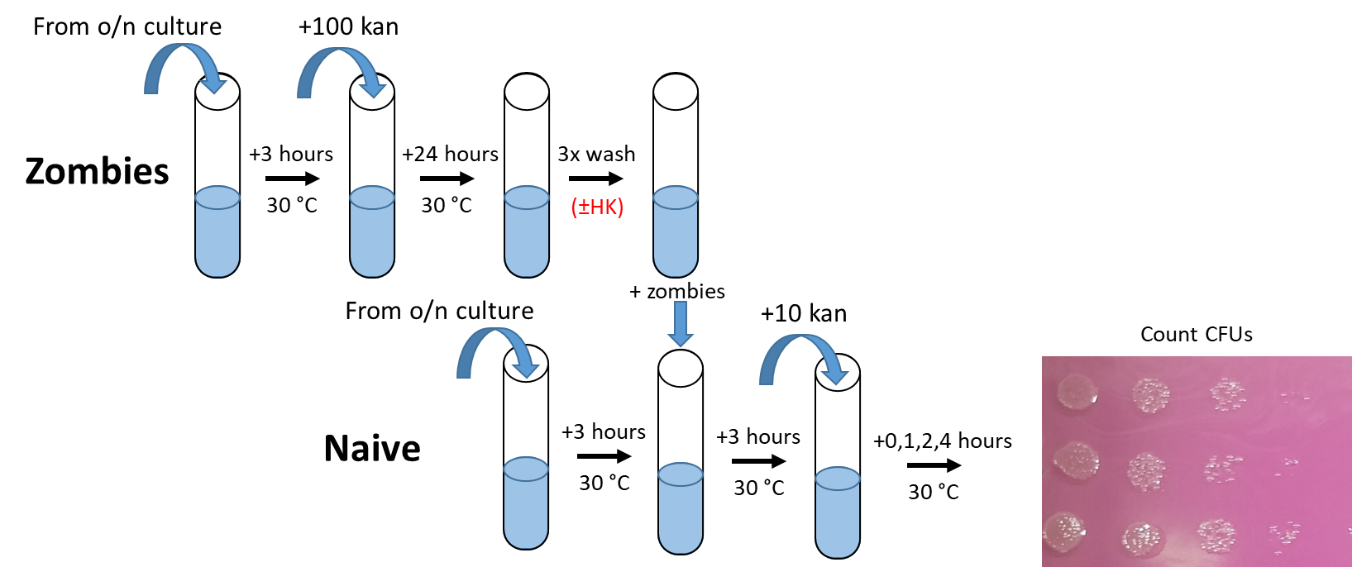

Schematic for the zombie protection assay. Zombies (top row): First cells are grown overnight (o/n) in LB, followed by dilution 1:100 into fresh PMM. After 3 hours in PMM, kanamycin is added and cells are incubated at 30 ºC for 18-24 hours. The next day, the zombies are washed 3x in fresh PMM (and heat killed if needed) and then resuspended in fresh PMM for 1 hour with no kanamycin. Naïve (bottom row): Cells are grown overnight in LB and diluted 1:100 in fresh PMM. After incubating for 3 hours at 30 °C, washed zombies are added in to the culture and the full cell suspension (zombies+naïve) is incubated for an additional 3 hours. Kanamycin is then added, and after the specified amount of treatment time, cells are used for a spot-plate CFU assay and counted the following day.

Figure S9:

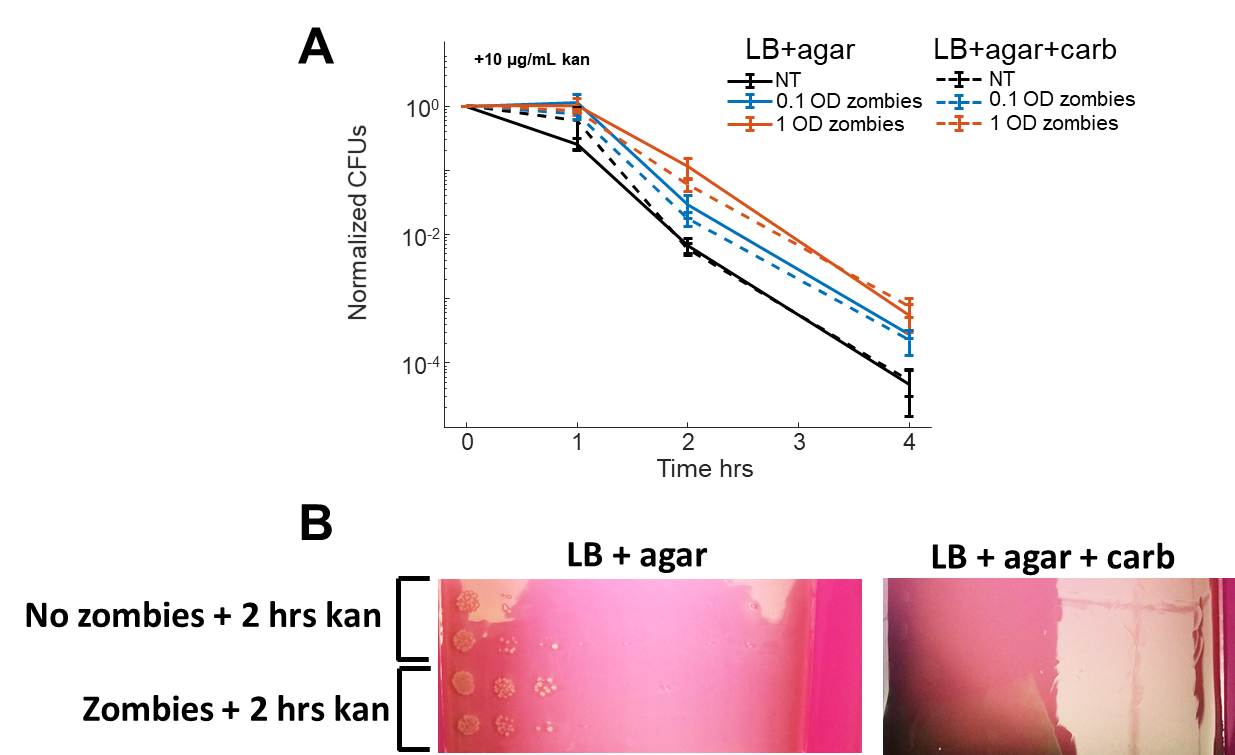

Observed protection of treatment naïve cells is not due to zombies re-entering the cell cycle. (A) Naïve cells were resistant to carbenicillin and zombies were susceptible. CFU curves of naïve cells with no-treatment (black), with 0.1 OD added zombies (blue), or 1 OD added zombies (red). The CFUs were plated on both LB+agar with no antibiotic (solid lines) or LB+agar+100 mg/mL carbenicillin (dashed lines). If zombies were contributing to CFUs, the carb plates would have lower CFUs since the zombies are not carb resistant. (B) Naïve cells were susceptible to carb and zombies were not. Images of CFU plates after two hours treatment with 10 μg/mL kanamycin. Though colonies grew on the LB+agar, there were none on the carb plate showing that after mixing with naïve cells there were no zombies that could form a colony.

Figure S10:

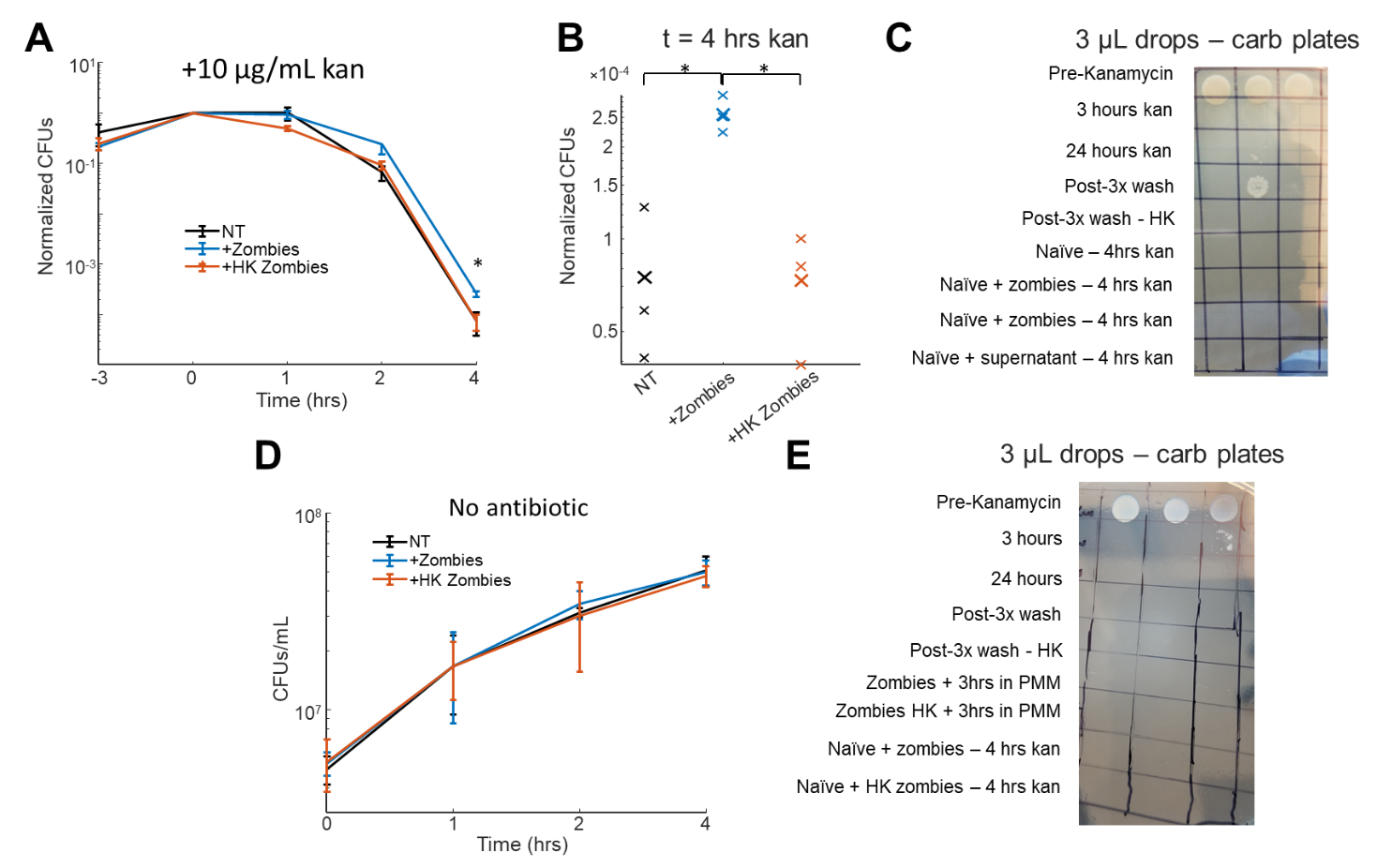

Actively metabolizing zombie cells are required for protection but do not affect growth rate of naïve cells. (A) CFU curves of naïve cells with no-treatment (black), with zombies (blue), or with heat-killed zombies (red). Heat killed zombies do not protect naïve cells, indicating metabolism of zombies is required. Error bars are the standard deviation of 3 biological replicates. (B) CFU measurements of 3 biological replicates after 4-hours kanamycin treatment. Each x indicates a biological replicate and the large X is the mean. ^*^ represents p < 0.05 in a two-sided t-test. (C) Spot plate with carb showing that no zombies can re-grow during the treatment with the naïve cells. Columns are 3 biological replicates. (D) CFU counts of naïve cells untreated (black), treated with zombies (blue), or treated with heat killed zombies (red) in the absence of antibiotics. There is no difference in growth rate between these conditions. (E) Spot plate showing that no zombie colonies contributed to the measurements in (D). Columns are 3 biological replicates.

Figure S11:

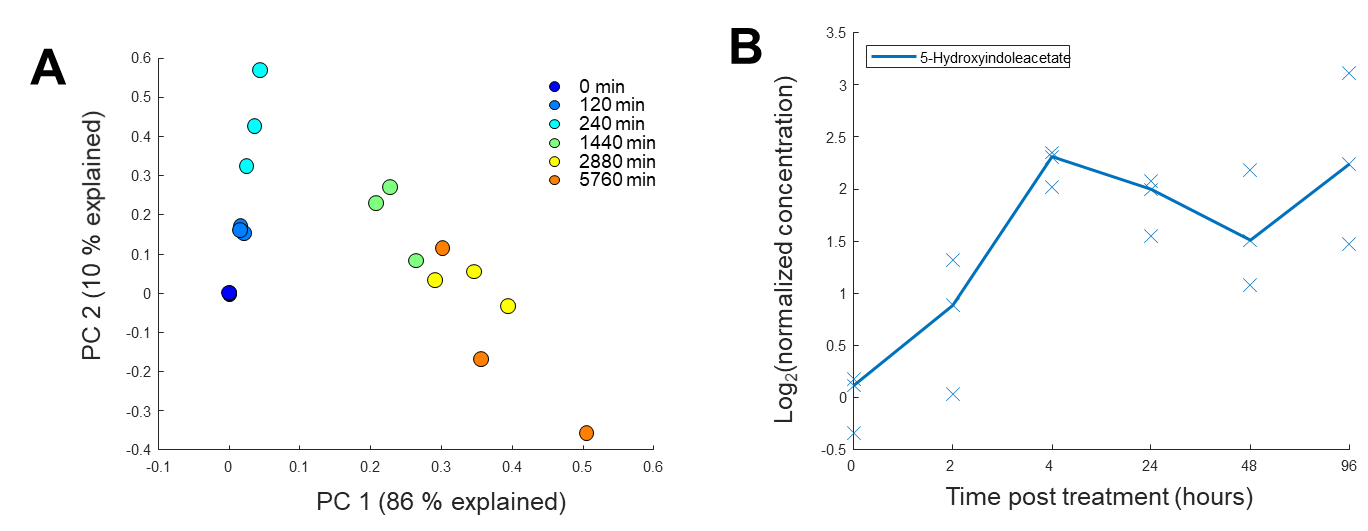

Metabolomics of zombie secretome shows increases in indole derivatives. (A) PCA decomposition of the metabolomic profile at each time point. The color represents the time point showing good clustering and changing profiles with time in aminoglycoside. The time is given in minutes post treatment. (B) Normalized change in supernatant concentration of 5-hydroxyindoleacetate. Each x is one biological replicate, and the line shows the change in the median at each time point.

Figure S12:

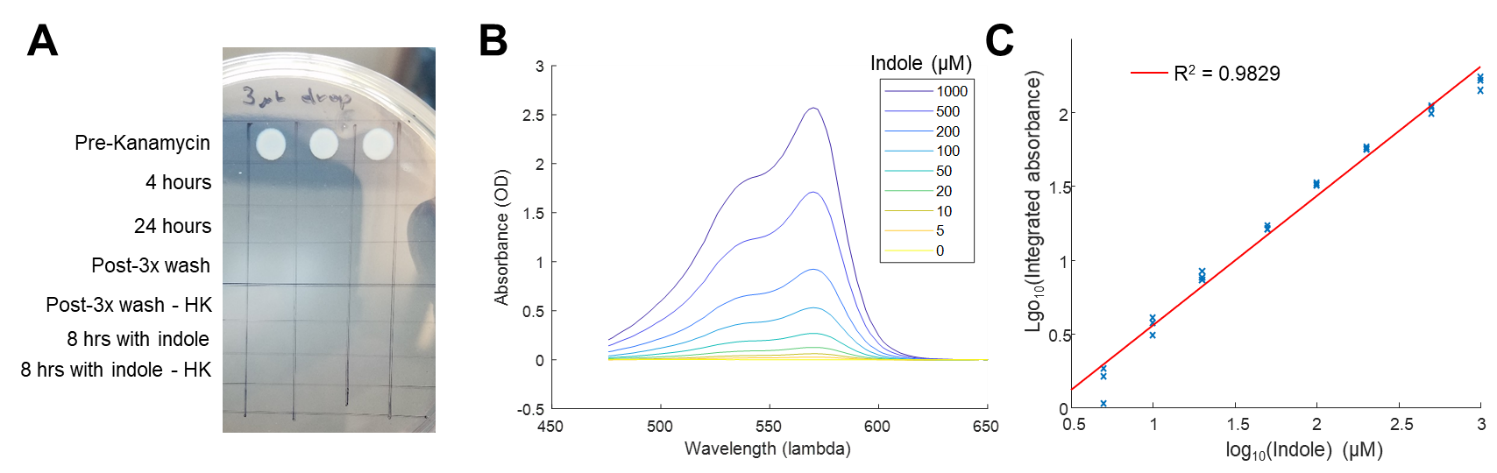

Measuring indole of the supernatant from zombies. (A) Spot plate showing that CFUs are below our detection limit (333 cells/mL) after treatment with kanamycin and when measuring indole (bottom two rows). (B) Absorbance spectra of Kovac’s reagent when mixed with a defined titration of indole. The concentration of indole is reflected by the color, and is given in μM. (C) Fitting the integrated intensity of the absorbance spectrum to a line shows good linear correspondence between absorbance and concentration. Each x is a replicate measurement, and the red line is a linear fit.
